# Functional Pathways of Biomolecules Retrieved from Single-particle Snapshots

**DOI:** 10.1101/291922

**Authors:** A. Dashti, M.S. Shekhar, D. Ben Hail, G. Mashayekhi, P. Schwander, A. des Georges, J. Frank, A. Singharoy, A. Ourmazd

**Affiliations:** Dept. of Physics, University of Wisconsin Milwaukee, 3135 N. Maryland Ave, Milwaukee, WI 53211; Beckman Institute for Advanced Science and Technology, and University of Illinois at Urbana-Champaign, Urbana, IL 61801, United States; Structural Biology Initiative, CUNY Advanced Science Research Center, City University of New York, New York, NY 10031; Dept. of Chemistry & Biochemistry, City College of New York, New York, NY 10031; Ph.D. Program in Biochemistry, The Graduate Center of the City University of New York, New York, NY 10016; Dept. of Biochemistry and Molecular Biophysics, Columbia University, 2-221 Black Building, 650 West 168^th^ Street, New York, NY 10032; Dept. of Biological Sciences, Columbia University, 600 Fairchild Center, New York, NY 10027; School of Molecular Sciences, Center for Applied Structural Discovery, Arizona State University, Tempe, AZ 85287, United States

**Keywords:** Conformations of biological molecules, molecular movies, allosteric regulation, ion channel, calcium release channel, ryanodine receptor, RyR, excitation contraction coupling, gating, cryo-electron microscopy, manifold embedding

## Abstract

We present a new approach to determining the conformational changes associated with biological function, and demonstrate its capabilities in the context of experimental single-particle cryo-EM snapshots of ryanodine receptor (RyR1), a Ca^2+^-channel involved in skeletal muscle excitation/contraction coupling. These results include the detailed conformational motions associated with functional paths including transitions between energy landscapes. The functional motions differ substantially from those inferred from discrete structures, shedding new light on the gating mechanism in RyR1. The differences include the conformationally active structural domains, the nature, sequence, and extent of conformational motions involved in function, and the way allosteric signals are transduced within and between domains. The approach is general, and applicable to a wide range of systems and processes.

## Introduction

In equilibrium, each conformational state of a macromolecule is occupied with a probability 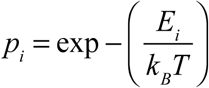, where *E*_*i*_ is the free energy of the conformational state *i*, *k*_*B*_ the Boltzmann constant, and *T* the temperature. Single-particle snapshots of a sufficiently large number of macromolecules will include all accessible conformational states, with the number of snapshots emanating from each state determined by its occupation probability. For a macromolecule with *N* atoms, the conformational landscape has 3*N* degrees of freedom.

Biological function, however, involves structural blocks, e.g., an alpha helix, or an entire molecular domain. Function can be thus described in terms of a small set of so-called *conformational coordinates*, each describing the concerted motions of a large number of atoms. The number of degrees of freedom exercised during unperturbed function, and the conformational coordinates relevant to such function must be experimentally determined. The choice of conformational coordinates is not unique, but independent, or at least mutually orthogonal coordinates are the most convenient.

Conformations can be represented as points on energy landscapes spanned by the conformational coordinates. The energy of each observed conformational state can be determined from the number of times it has been sighted, through the Boltzmann relation between the occupation probability *p*_*i*_ and energy of the state *E*_*i*_. Function unfolds along heavily populated (“least-action”) paths on energy landscapes (*1*).

It has long been recognized that landscapes specifying the free energy of each conformation offer a powerful framework for discussing function (*2*, *3*), and the literature abounds with sketches of such landscapes. The few experimentally determined energy landscapes, however, are predominantly one-dimensional (i.e., involve a single conformational coordinate), and are described in terms of qualitative, ad hoc, or externally imposed reaction coordinates not guaranteed to capture the changes relevant to function.

In the absence of suitable, experimentally determined energy landscapes, efforts to infer function often rely on powerful maximum likelihood classification methods (*4*, *5*) to sort single-particle snapshots into a user-defined number of discrete conformational clusters. A 3D structure is then extracted from each cluster. In general, the sequence in which these structures appear in the course of function, and indeed their relevance to function are unknown. Under such circumstances, functional inference necessarily involves linear interpolations between clusters, if only conceptually. As the number of ways in which two discrete structures can be transformed into each other is essentially unlimited, functional inference by discrete clustering is fraught with difficulty.

The primary goals of this article are as follows. First, to demonstrate the compilation of experimental energy landscapes associated with complex biological function, including those involving more than one landscape. Second, to identify the conformational paths associated with function, even when they involve interlandscape transitions. Third, to demonstrate the motions revealed by such functional analysis are significantly different from those inferred by discrete clustering methods. And finally, to outline the new biological insights gained by studying the continuous conformational changes associated with function.

For concreteness, the comparison is made with reference to ryanodine receptor type 1 (RyR1), a large, functionally complex, fourfold-symmetric (C4) ion channel in the sarcoplasmic reticulum membrane. RyR1 is a calcium-activated calcium channel critical to excitation/contraction coupling in skeletal muscle. Several recent cryo-EM studies have characterized, in exquisite detail, the many discrete structures obtained by clustering techniques (see, e.g., (*6*-*10*)). The functional information inferred from these studies includes the conformational states assumed by the channel (*6*-*8*), and the effects of activation and gating induced by ligand binding (*9*, *10*). Nonetheless, our understanding of key functional processes, such as the allosteric coupling between the cytoplasmic shell and pore of the channel, remains incomplete. RyR1 thus offers an ideal opportunity to compare the conformational information deduced by standard clustering methods with that revealed by the functional approach used here.

This article is organized as follows. We first outline how manifold-based geometric machine learning (*1, 11-14*) can be used to determine the experimental energy landscapes of RyR1 with and without ligands in terms of rigorously derived, mutually orthogonal conformational reaction coordinates associated with ligand binding. This provides important insights into the mechanisms underlying the process of ligand binding, including the presence of multiple routes to ligand binding, and the associated functional motions. These observations are then contrasted with the results obtained by analyzing discrete structures. The comparison reveals major differences in the delineation of functionally active structural domains, the nature, sequence, and extent of motions associated with function, and the way allosteric signal is propagated to functionally important remote sites. The article concludes with a discussion of the new biological insights provided by the new approach.

### Functional analysis of RyR1

We have previously demonstrated that experimental single-particle snapshots of molecular machines idling in equilibrium on a single energy landscape can be used to determine functionally relevant conformational motions in terms of rigorously derived orthogonal coordinates (*1*). Such an energy landscape reveals all conformations with energies up to an upper limit set by the vanishing occupation probability of high-energy states. The key point is that thermal fluctuations in equilibrium lead to sightings of all states up to the limit set by the number of snapshots in the dataset (SM section 1).

An important feature of the present study is the pooling of cryo-EM snapshots from two experiments. In one experiment, RyR1 macromolecules were in equilibrium with a thermal bath without any activating ligands. In the other, the macromolecules were in equilibrium with both a thermal bath and a reservoir of ligands, specifically calcium, ATP, and caffeine (*9*) (SM section 2). This pooling of data allows both species (with and without ligands, henceforth ±ligand) to be described in terms of the same set of mutually orthogonal conformational coordinates. The resulting ±ligand energy landscapes reveal the heavily populated conformational conduits, which we associate with routes relevant to ligand binding (Fig. 1A).

**Fig. 1:**
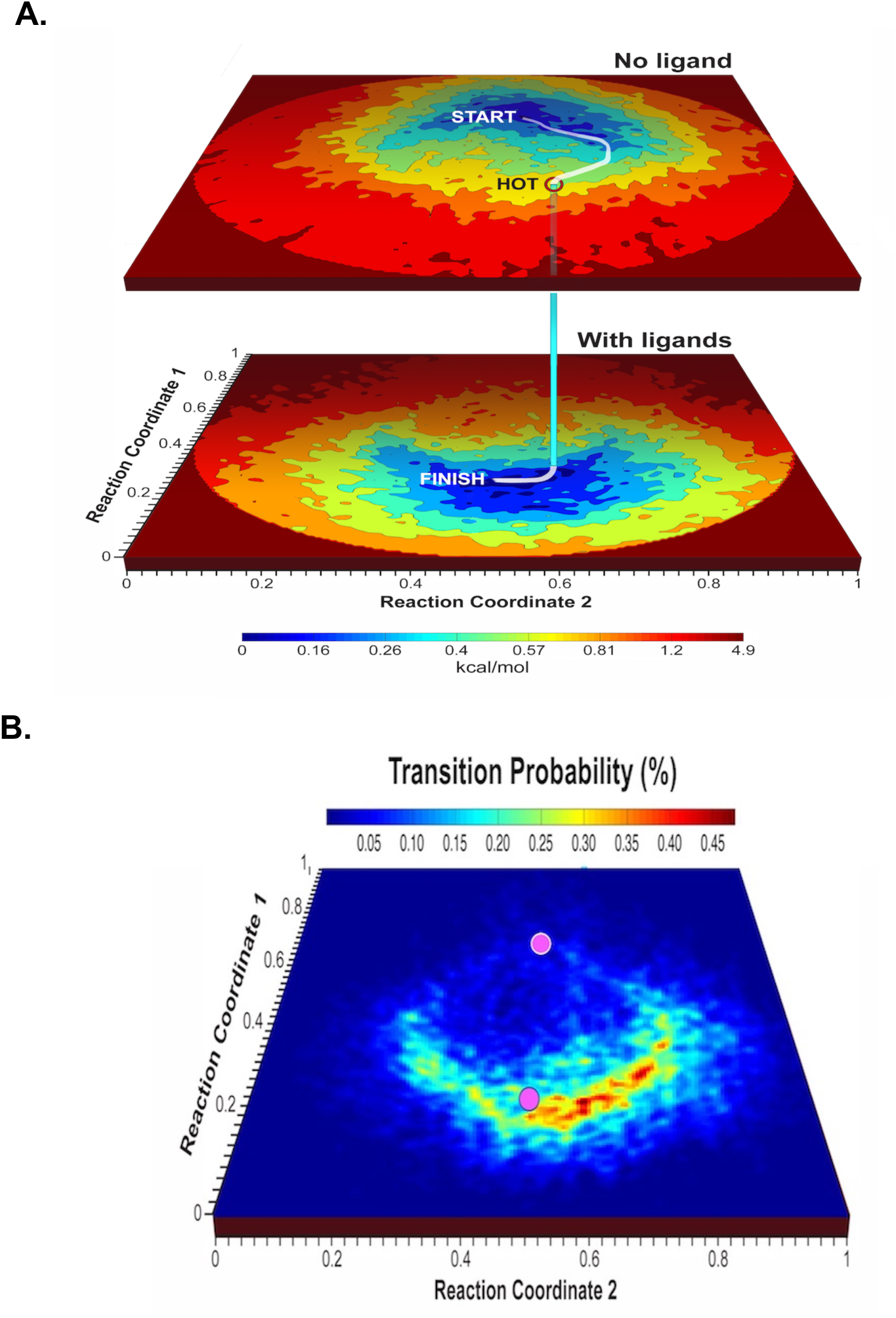
Energy landscapes for RyR1 with and without ligands, and map of the probability of a transition from the no-ligands to the with-ligands landscape. **(A)** Energy landscapes without ligands (upper surface), and with ligands (Ca^2+^, ATP, and caffeine) (lower surface). The landscapes are described in terms of the most important two of a common set of orthogonal conformational coordinates. The curved path represents a high-probability route to the binding of ligands. This path starts at the minimum-energy conformation of RyR1 without ligands (“START”), follows the conduit of lowest energy to a point with a high probability of transition to the with-ligands energy landscape (“HOT”), and terminates at the minimum-energy conformation with ligands (“FINISH”). **(B)** Probability map for transitions from the energy landscape without ligands to the energy landscape with ligands. The axes are the same as in Fig. 1A, with magenta discs indicating the positions of the minima of the energy landscapes at (0.7, 0.57) and (0.23, 0.56), respectively.

Subject to reasonable assumptions, Fermi’s Golden Rule (*15*, *16*) (SM section 3) is then used to estimate the transition probability between the two landscapes, with “hotspots” identifying the most probable transition points between the landscapes (Fig. 1B). Three – dimensional (3D) movies compiled along heavily populated conduits on these landscapes reveal the conformational motions associated with ligand binding, in some regions with near-atomic resolution.

In greater detail, the 791,956 cryo-EM snapshots of RyR1 molecules analyzed in this study comprised about the same number of molecules in equilibrium with reservoirs with and without ligands (Ca^2+^, ATP, and caffeine) prior to cryo-freezing (*9*). (For details see SM section 2.) For a discussion of the residual ligand concentration in the no-ligand solution, see SM section 2.) These snapshots were grouped into 1,117 uniformly spaced orientational bins by standard procedures (*5*). Geometric (manifold-based) analysis of the pooled dataset revealed two significant conformational reaction coordinates, each describing a concerted set of continuous changes (RC1 and RC2 for short).

The architecture of the ryanodine receptor can be divided into three major regions: the channel pore, responsible for calcium efflux from the sarcoplasmic reticulum; an activation core, responsible for ligand binding and channel activation; and a large cytoplasmic shell serving as a platform for the binding of many regulatory proteins. Broadly speaking, conformational changes along RC1 involve the shell, the activation core, and the pore; those along RC2 the shell (SM section 4).

Fig. 1A shows the ±ligand energy landscapes of RyR1. Assuming the probability of a collision (not binding) with a ligand is independent of conformation, the probability of a transition between equivalent points on the two landscapes can be estimated from Fermi’s Golden Rule for the period immediately after the exposure of RyR1 molecules to the reservoir containing ligands (SM section 3). As shown in Fig. 1B, the inter-landscape transition probability displays specific “hotspot” regions, where a significant number of ligand-free and ligand-bound macromolecules have the same conformation. The most probable routes to ligand binding start from the region of lowest energy on the –ligand landscape (“START” in Fig. 1A), reach one of the hotspot transition points (“HOT”) with a probability of ~2%, cross to the +ligand landscape with ~ 0.45% of the probability of a collision with a ligand, and terminate in the region of lowest energy on the +ligand landscape (“FINISH”). This means ~ 0.01% of collisions with a ligand lead to binding.

The displacement of inter-landscape transition hotspots (red regions in Fig. 1B) from minimum energy regions (magenta dots) on both landscapes highlights the need for significant conformational changes before *and* after transition between the two energy landscapes. At the same time, the presence of several inter-landscape transition hotspots reveals a multiplicity of routes to ligand binding with comparable transition probabilities. The curved nature of all routes to binding (e.g., the white line in Fig. 1A) emphasizes the inappropriateness of deducing functional information from discrete “START” and “FINISH” structures at the extremes of the conformational range.

The results outlined above already elucidate longstanding questions regarding the “population shift” vs. “induced fit” models of ligand binding. Broadly speaking, “population shift” (*17*) requires a conformational change *before*, “induced fit” (*18*) a conformational change *after* ligand binding. Our results show ligand binding, at least in RyR1, proceeds via a continuum of conformations increasingly different from, and at higher energies than, the minimum-energy conformation on the –ligand landscape. These higher-energy conformations are reached thermally via “population shift”. Collision with a ligand then transfers RyR1 to the +ligand energy landscape, where a downward slope in energy drives further continuous conformational changes to the minimum-energy, ligand-bound state. The conformational changes *after* collision with a ligand constitute an “induced fit”. Our results thus show that each of the two opposing models describes a different part of the actual process; at least for RyR1, binding entails specific conformational changes both before *and* after collision with a ligand.

The exact apportionment of the conformational changes before and after collision depends, of course, on the point at which the transition to the +ligand landscapes takes place. Although transitions can occur over a relatively broad region, the highest probabilities are concentrated at a few “hotspots” (Fig. 1B). The positions of the ±ligand energy minima relative to the broad region of significant transition probability indicate that most ligand-binding events in RyR1 involve a greater element of “population shift” than “induced fit”, as suggested in (*19*), but this could be system-specific.

### Functional analysis versus linear interpolation between discrete structures

The above discussion makes it clear that our functional analysis provides a wealth of new information on the mechanism of ligand binding. The new information includes the conformational coordinates relevant to function, the functional routes to ligand binding, the points at which transitions occur between the +ligand and –ligand landscapes, and the transition probabilities and branching ratios for different functional paths. None of this information can be obtained by studying discrete clusters.

Indeed, the availability of energy landscapes enables us, for the first time, to investigate the relationship between the discrete classes produced by maximum likelihood clustering (*20*) of the same RyR1 snapshots (*7*, *9*). The data analyzed here were previously clustered into 16 conformational classes, two of which were classified as “junk” (*9*). The distribution of the snapshots (on the energy landscapes) contributing to each class is shown in Fig. 2. It is difficult to discern a systematic relationship between the positions of the different classes. Indeed, many of the clusters do not correspond to notable regions of the landscapes, with snapshots from “junk” clusters randomly distributed over the landscapes. Class 2 (no ligands) and class 3 (with ligands) were taken in the previous cluster-based analysis (*7*, *9*) to represent the functionally relevant extremes of the conformational range. Function was then inferred by interpolation between the 3D structures obtained from these two classes. Below, we compare the conformational changes obtained from such interpolation, with the changes associated with the functional trajectory on the energy landscapes (Fig. 1).

**Fig. 2:**
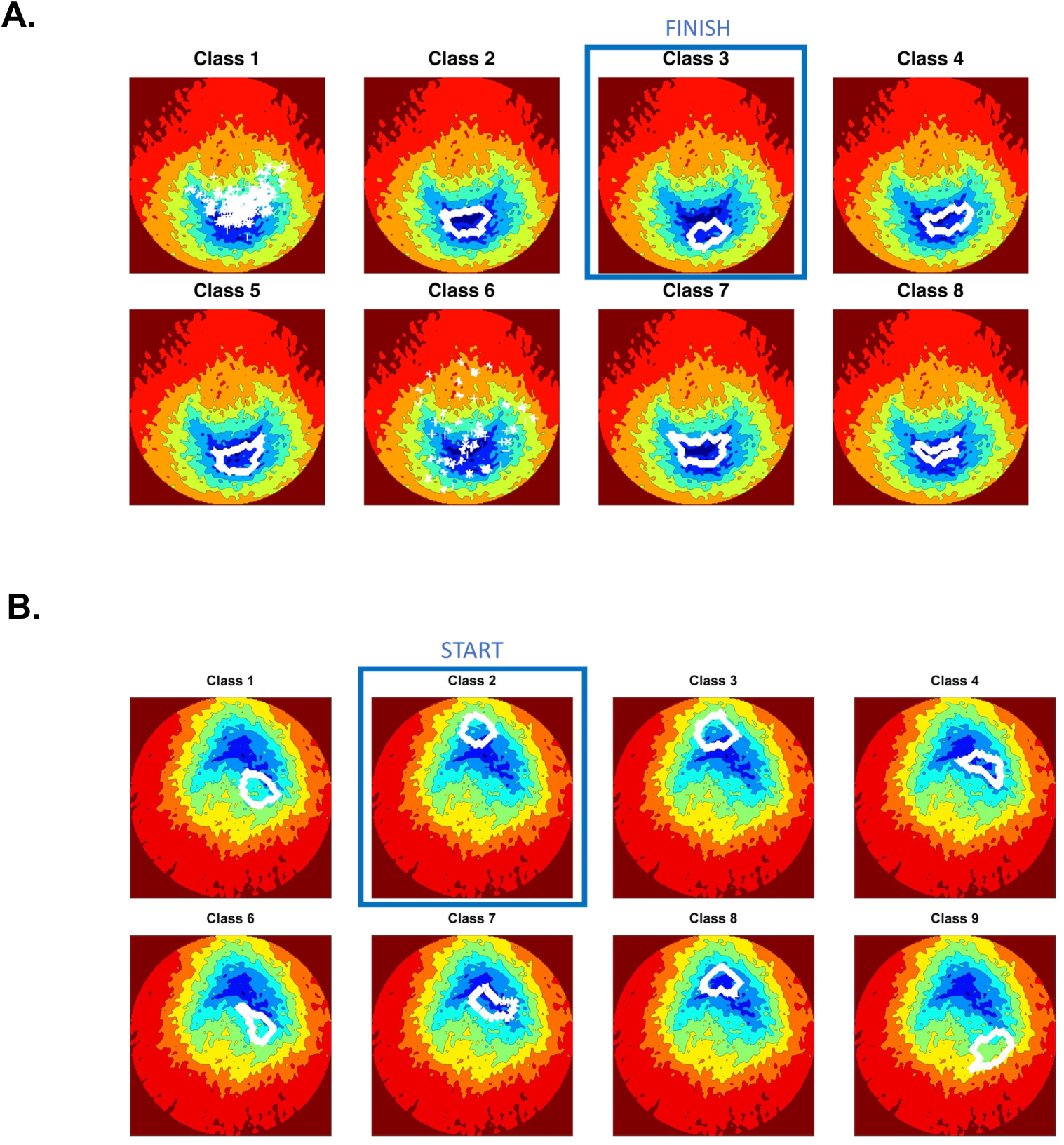
Positions of discrete conformational clusters obtained by maximum-likelihood analysis on the energy landscapes of Fig. 1A. **(A)** No ligands. **(B)** With ligands. The boxed classes were previously identified as representing the extremes of the conformational range, and used to infer function by discrete cluster analysis (*9*).

Before doing so, however, we note two important points. First, the extent to which maximum likelihood clustering is based on functionally relevant conformational reaction coordinates is unknown. The position of a discrete cluster on the relevant energy landscape (Fig. 2) thus represents a *projection* from an unknown space onto the space spanned by the two conformational reaction coordinates most relevant to ligand binding and channel gating. Second, interpolation (“morphing”) between two or more static structures along a putative functional path is generally acknowledged as invalid. But the approach is nonetheless often used to infer function from static structures, because the functional path connecting two discrete clusters is unknown. In the absence of other information, this makes functional inference by interpolation all but inevitable, if only conceptually.

We now turn to the conformationally active structural domains and their motions. Fig. 3 and Movies 1 and 2 compare the displacements revealed by our functional analysis with those inferred from interpolating between the discrete clusters (ringed classes in Fig. 2) used in a previous study (*10*). These discrete classes lie close to the energy minima on the ±ligand energy landscapes, i.e., the START and FINISH of the inter-landscape functional path.

**Fig. 3:**
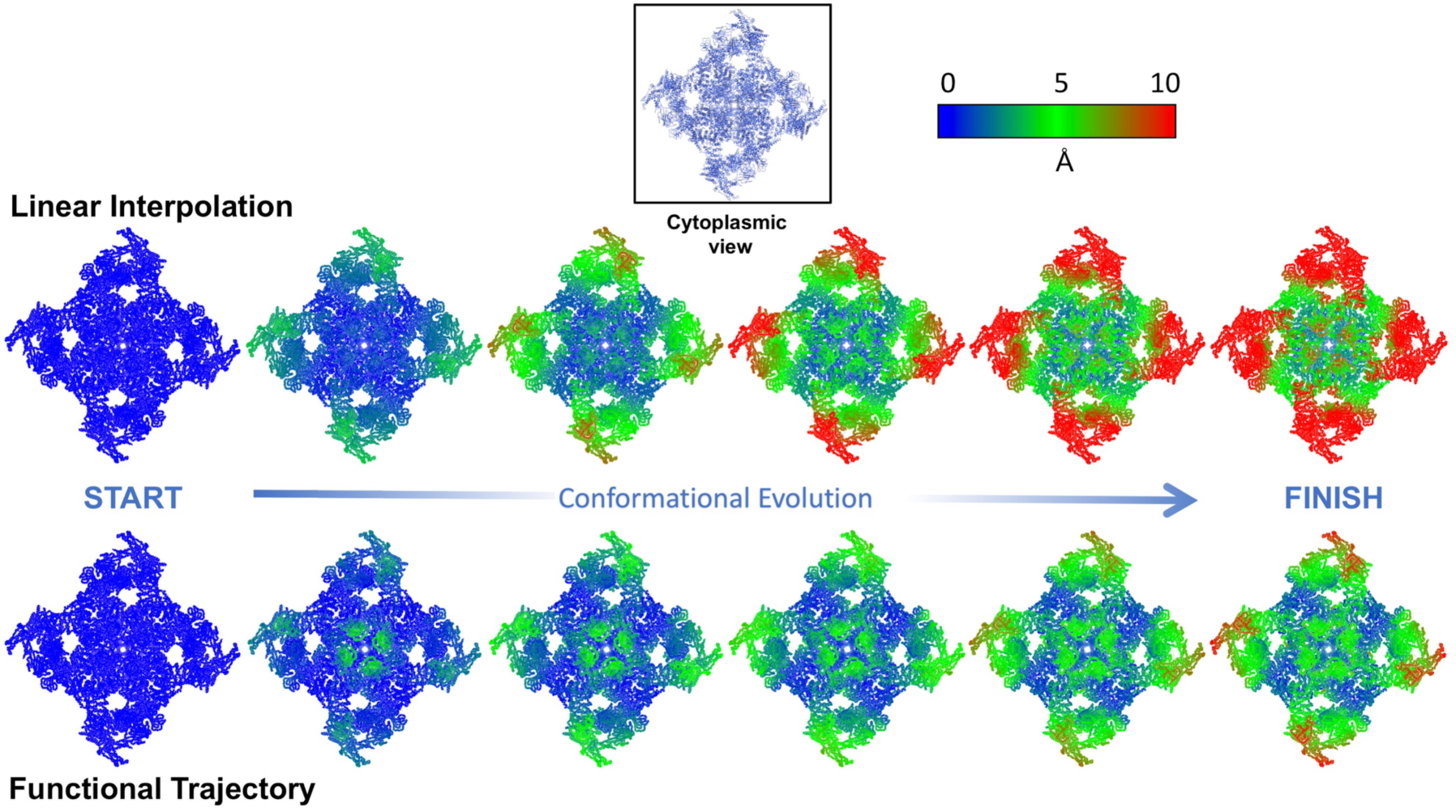
**Overview of the conformational evolution along the functional route of Fig. 1A with the trajectory vs. linear interpolation between the two maximum-likelihood classes previously used for functional inference** (*9*) (boxed in Fig 2). The RyR1 atomic model is shown in sphere representation, with color indicating the displacement of each atom from its “START” position (see color bar).

There are major differences between the results of functional analysis and those inferred by interpolating between the discrete clusters. These differences include the structural domains involved in motion, as well as the sequence and extent of displacements (Figs. 3-5, Movies 1-6). For example, in contrast to the results from cluster analysis, functional analysis shows that: a) the N-terminal domains (NTD) lead the sequence of motions; b) a significant section of the macromolecule remains rigidly static during function; and, c) the motions in the activation core and shell are coupled (Fig. 3, Movies 1-2).

The above conformational changes are substantially different from those inferred by interpolating between the discrete structures, even though the clusters used are close to the termini of the functional path. Conformational changes specific to the N-terminal domains have not been described in previous high-resolution cryo-EM studies, nor specific elements of the shell been shown to be more rigid than others (*8, 9, 21*).

Importantly, functional analysis reveals the motions involved in pore opening are significantly different from those inferred by discrete cluster analysis. Specifically, functional analysis shows the atomic displacements in the pore domain during channel opening are primarily located in the *transmembrane* region of the channel/pore scaffold (Fig. 4 and Movies 3-4). RyR1 function is known to be sensitive to mutations in this region; 40% of such mutations are implicated in Central Core Disease (see, e.g., http://www.uniprot.org/uniprot/P21817). In contrast, the displacements inferred from discrete cluster analysis primarily involve the *cytoplasmic* region of the pore-lining helix S6. These results graphically highlight the limitations of using static clusters for functional inference.

**Fig. 4:**
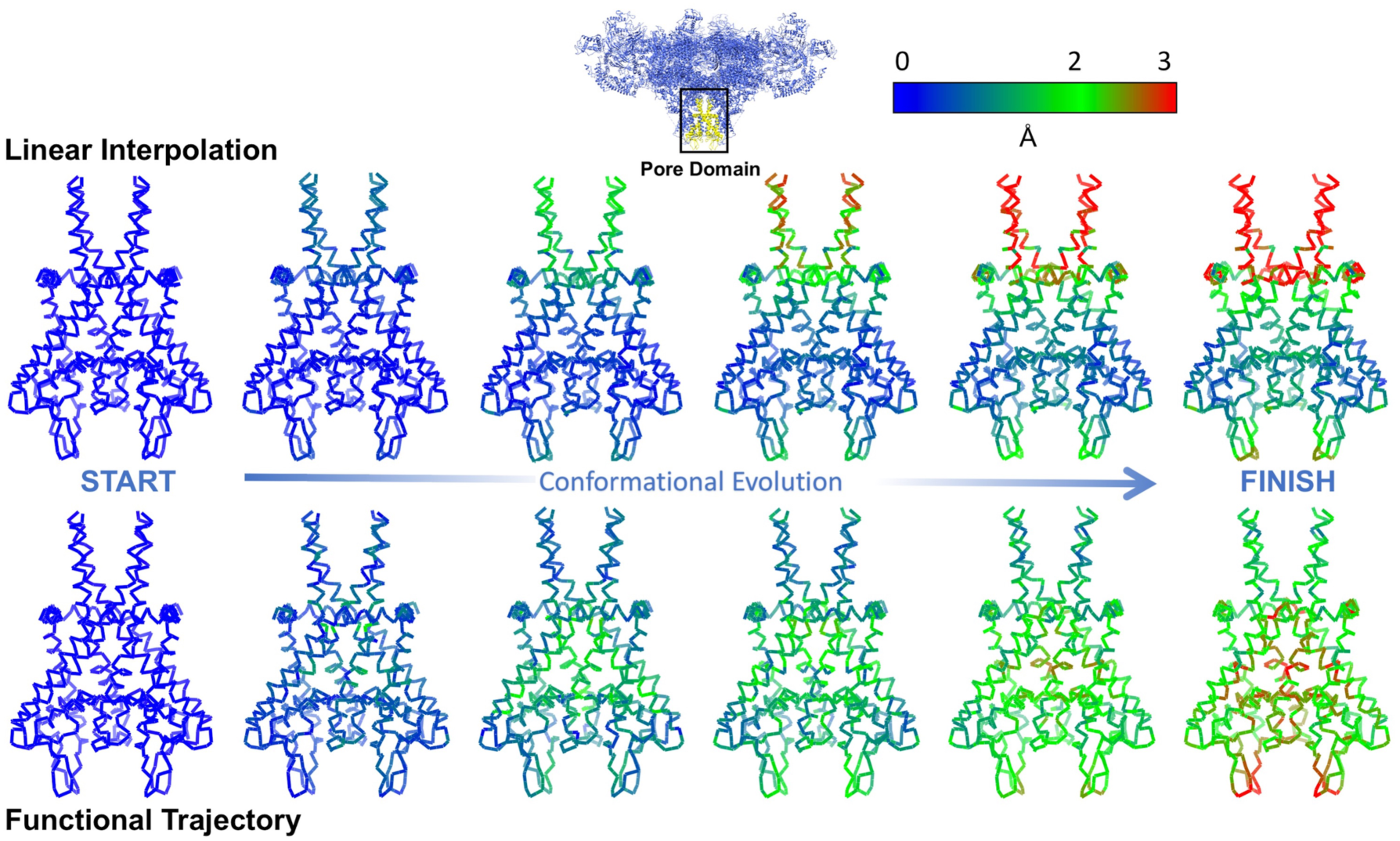
**Conformational evolution in the pore domain (residues: 4820-4956) along the functional route of Fig. 1(A) vs. the changes inferred by linear interpolation between classes previously used for functional inference** (*9*) (boxed in Fig. 2). The RyR1 carbon-alpha backbone is shown in stick representation, with color indicating the displacement of each atom from its “START” position (see color bar). The functional trajectory shows the largest conformational changes (red) occur in the transmembrane region of RyR1. In contrast, the largest changes obtained by interpolation between discrete clusters are located in the cytoplasmic region of the molecule.

### Insights into the RyR1 gating mechanism

We now discuss the implications of our functional results for the allosteric mechanisms responsible for channel pore opening upon binding of activating ligands. This discussion is illuminated by distance measurements at important sites, which quantify the consequences of functional motions.

The atomic coordinates of RyR1 obtained by modeling along the functional trajectory reveal the conformational changes at the binding sites of the ligands Ca^2+^, ATP and caffeine as ligand binding proceeds. Starting at the minimum-energy point of the –ligand landscape, the Ca^2+^ and ATP binding sites gradually contract until the transition to the +ligand landscape, after which the conformations of the binding-sites stabilize (Fig. 6). This is in line with the “population shift” view described earlier. The caffeine binding site, on the other hand, displays a more complex behavior: two aromatic amino-acids approach each other to accommodate, and potentially stabilize, caffeine binding, while two orthogonal amino-acids move apart (Fig. 7). The upward movement of the backbone supporting Ile 4996 may explain the calcium potentiation effect of caffeine (*22*), as this would bring the calcium-coordinating amino-acid Thr 5001 on the opposite side of this loop closer to its calcium-bound position (Fig. 6).

**Fig. 5:**
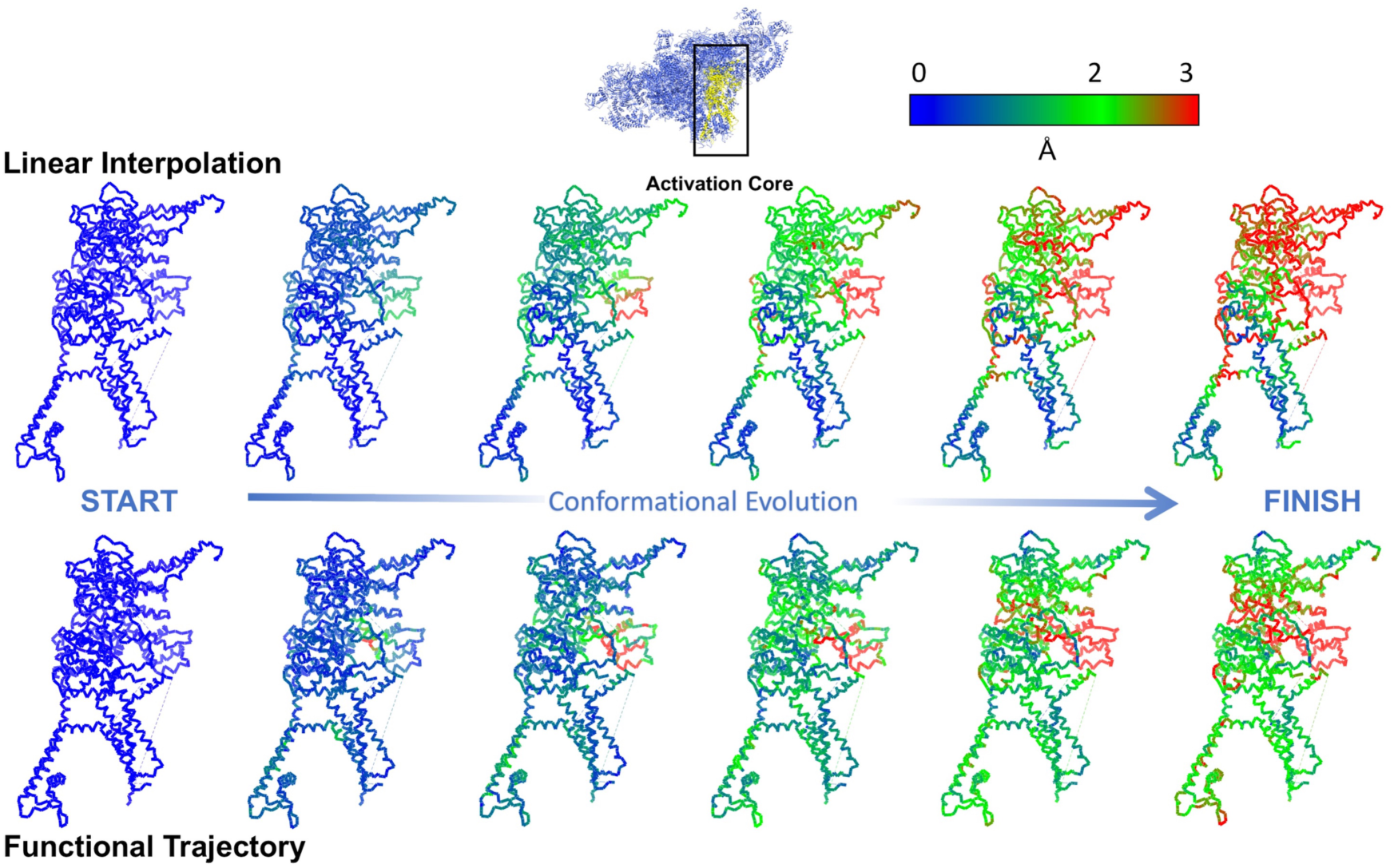
**Conformational evolution in the activation core domain (residues: 3614-5037) along the functional route of Fig. 1(A) vs. the changes inferred by linear interpolation between classes previously used for functional inference** (*9*). The RyR1 backbone structure is shown as sticks, with the color of each atom indicating its atomic displacement (see color bar). The largest atomic displacements along the functional trajectory are confined to a narrow allosteric conduit connecting the ligand binding sites to the EF-hand. In contract, interpolation between discrete clusters results in conformational changes distributed over a large part of the molecule.

**Fig. 6:**
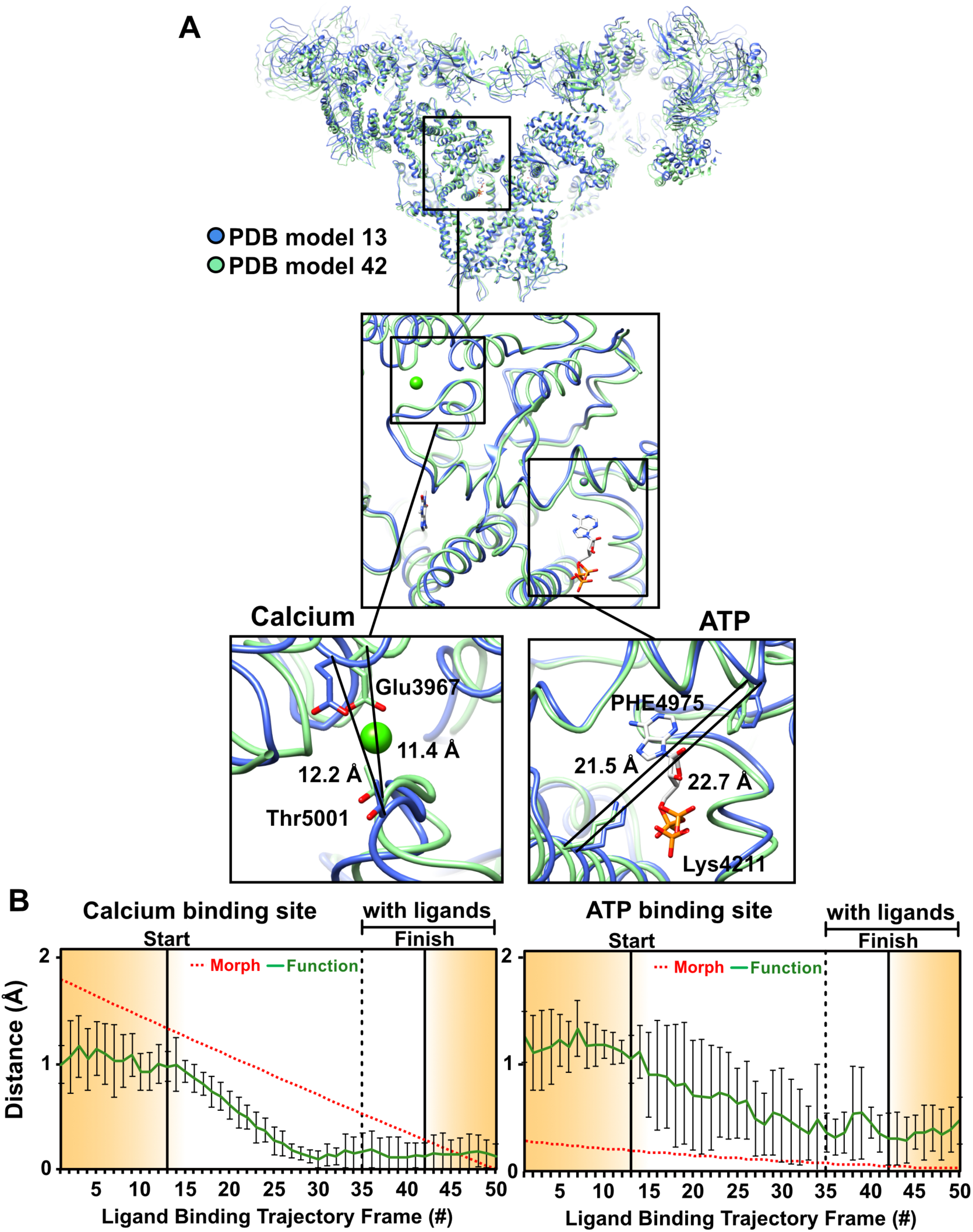
**Conformational changes at RyR1 ligand binding sites along the functional route of Fig. 1(A)**, augmented with excursions from the minimum-energy points along RC1. These excursions are in the increasing RC1 direction on the no-ligand landscape, and in the decreasing RC1 direction on the with-ligand landscape. **(A)** The general region, and the specific sites examined in detail. Distance variations between two amino acid backbones on opposite side of the ATP and calcium are shown. The amino acids used for measurement are represented as sticks, the rest of the molecule in blue ribbon for the model corresponding to “START” in Fig. 1(A), and in green ribbon for model corresponding to “FINISH”. **(B)** Distance variations between carbon-alpha backbone atoms of two opposing residues at each of the calcium and ATP binding sites for 50 structures along the functional and interpolated trajectories. For the latter, 50 structures were extracted from a morph between a model of RyR1 without ligands (PDB: 5TB4) and a model of RyR1 with calcium, ATP, and caffeine (PDB: 5T9V). To analyze the functional route, six published models (PDB: 5TB4, 5T9R, 5TAP, 5T9V, 5TAL, 5TAQ) were each fitted and refined into the maps along the trajectory. Error bars represent the full scatter (not standard deviation) of the results obtained with different starting models.

**Fig. 7:**
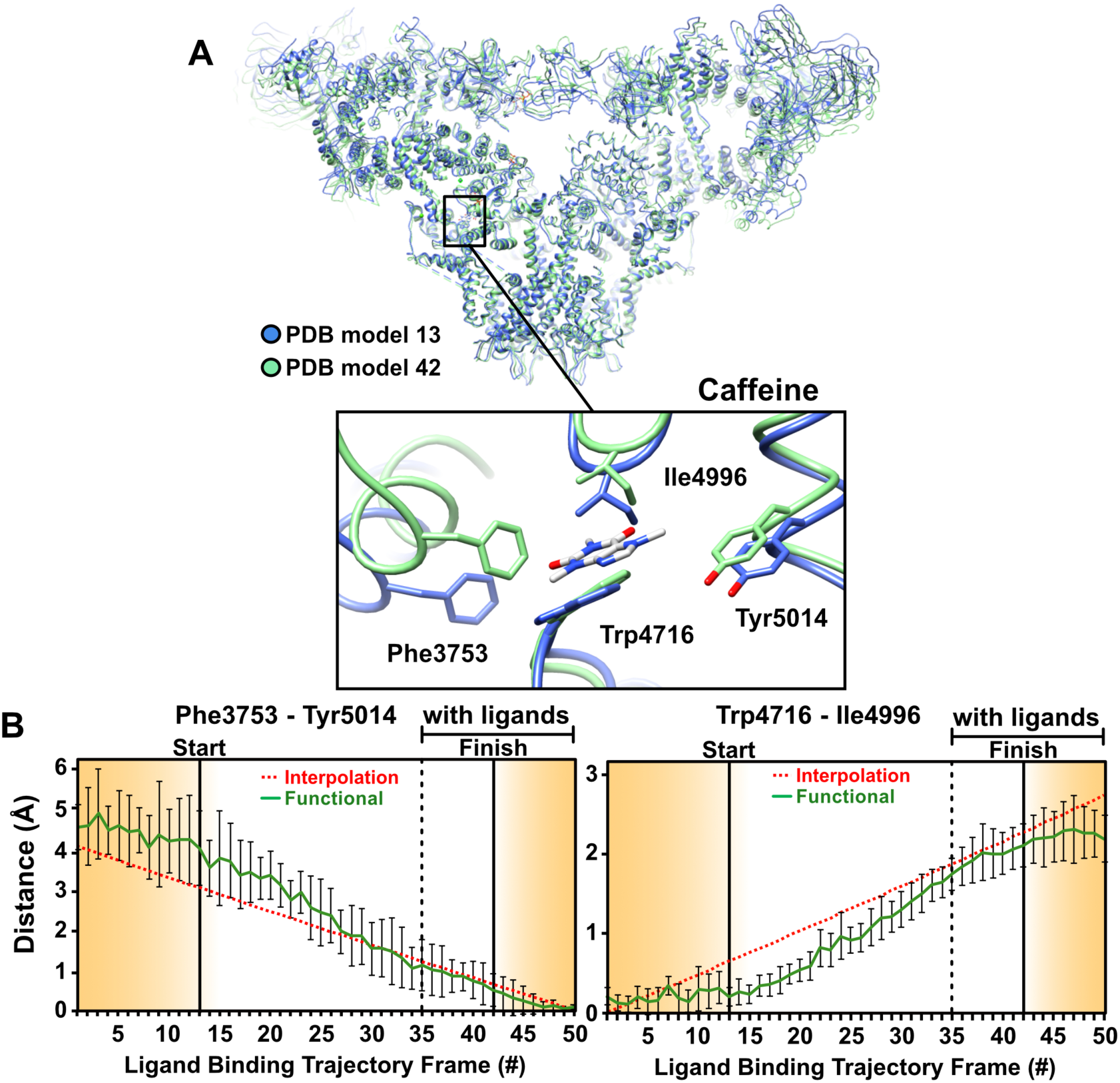
**RyR1 conformational changes at the caffeine binding site along the functional trajectory of Fig. 1(A)**, augmented with excursions from the minimum-energy points along RC1. These excursions are in the increasing RC1 direction on the no-ligand landscape, and in the decreasing RC1 direction on the with-ligand landscape. **(A)** The general region, and the specific site (inset) examined in detail. The molecule is shown in blue ribbon for the model corresponding to “START”, and in green ribbon for model corresponding to “FINISH” in Fig. 1(A). **(B)** Distance variations between two pairs of amino acid backbones on opposite side of caffeine. The amino acids used for measurement are represented as sticks.

It is known that ligand binding stabilizes the activation domain in a conformation suitable for pore opening (*9*). Our results show this activated state can be assumed also in the ligand-free state, albeit with low, or no conductance (Fig. 8). Consistent with the low probability of channel opening in the absence of calcium and ATP (23), the energy landscape shows only ~2% of RyR molecules assume the activated conformation in the ligand-free state. Further pore opening requires the binding of a ligand followed by an “induced fit” to the minimum-energy conformation of the ligand-bound receptor. The relatively broad +ligand energy minimum (Fig. 1) is also consistent with previous observations: Brownian motions of the shell (*6*-*8*) give rise to conformational dispersion along RC2; and the spikes in conductance, interpreted as channels flickering between the open and closed states (*23*), cause dispersion along RC1.

**Fig. 8:**
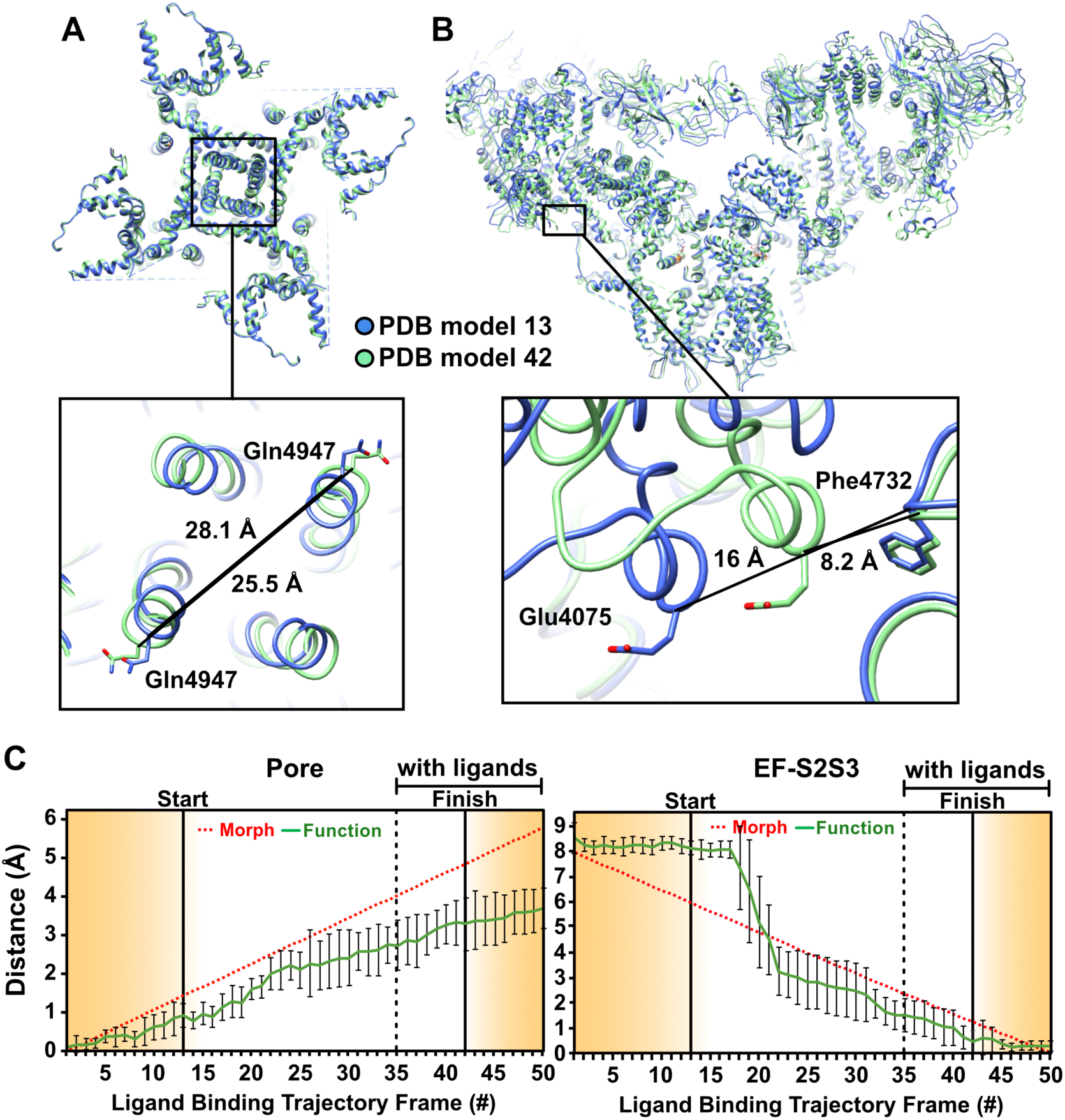
**Conformational changes in the pore and EF-hand domains along the functional trajectory**, augmented with excursions from the minimum-energy points along the conformational coordinate RC1. The general region, and the specific sites (insets) examined in detail. Distance variations between two amino acid backbones: **(A)** In the pore measured at Gln4947. **(B)** Between EF-hand (EF) at Glu4075 and S2S3 domain at Phe4732. The amino acids used for measurement are represented as sticks, the rest of the molecule in blue ribbon for the model corresponding to “START” in Fig. 1(A), and in green ribbon for model corresponding to “FINISH”. **(A)** Between carbon-alpha backbone atoms of two opposing residues. The measurements performed and plotted as described in Fig. 6.

Despite extensive work (see, e.g., (*9*)) it has proved difficult to clarify the way the conformational changes associated with ligand binding in the activation domain lead to gating and pore opening. Our distance measurements reveal potentially important atomic motions as the functional trajectory is traversed, most strikingly along a previously unobserved allosteric conduit connecting the ligand-binding sites in the Csol domain to the EF-hand (Fig. 5, Movies 5-6). Frame-by-frame measurements indicate the displacements begin at the calcium binding site, propagating along a narrow “vein” to the EF-hand (Fig. 5). Movement of the EF-hand has been observed before (*8*, *9*). Our analysis, however, uncovers a previously unobserved allosteric conduit between the ligands binding sites and the EF-hand, revealing the mechanical motions underlying signal transduction (Movie 5). In contrast, the displacements inferred from discrete cluster analysis are distributed uniformly over large regions, with no special feature indicating targeted signal transduction (Fig. 5, Movie 6). The functional analysis shows the narrow band of displacements first appears on the –ligand landscape, i.e., before ligand binding. This further supports the notion that “population-shift” is involved in the first part of the ligand-binding process, whereby a ligand stabilizes fleeting conformational fluctuations present before ligand binding.

The observed coupling of the EF-hand movement to ligand-binding points to the potential role of the EF-hand in gating, or its regulation. The movement leads to an interaction between the EF-hand and the S2S3 domain of the pore pseudo-voltage sensor. The small movement of the S2S3 domain associated with that interaction may be relevant to gating, but this hypothesis remains to be tested.

As noted earlier, the NTD is among the first domains affected by the transition between ligand-free and ligand-bound states (Fig. 3, Movies 1-2), suggesting a potentially important role for NTDs in gating. Indeed, the NTDs give rise to important interprotomer interactions, which are lost during channel dilation and subsequent pore opening, and a number of disease causing mutations are located at these interfaces (*24,25, 26*). It is thus important to understand whether NTDs and other inter-domain contacts involved in the gating mechanism are destabilized by the binding of ligands prior to pore opening, or they are sufficiently weak to be broken by Brownian motions during pore opening, a mechanism previously described as the “zipper hypothesis” (*27*). To clarify this question, we investigated the distance between interprotomer contacts as the functional path is traversed. The analysis was limited to backbone-to-backbone distances, as the resolution of our present study is currently limited by the number of snapshots to ~ 4.5 Å in the core of the channel, which precludes reliable measurement of side-chain positions (SM section 5).

Two interprotomer contacts display significantly nonlinear behavior, suggesting a possible role in gating (Fig. 9). The first such contact is formed between the EF-hand and the S2S3 domain of the neighboring protomer, as outlined above. We observe a stepwise motion bringing these two domains into close proximity well before the transition to the ligand-bound state and pore opening (Fig. 9B). The EF-hand movement is therefore not correlated with pore opening, but with channel activation. As deletion of the EF-hand does not affect channel activation (*28*), a role in regulation, or calcium inactivation appears more likely (*29*).

**Fig. 9:**
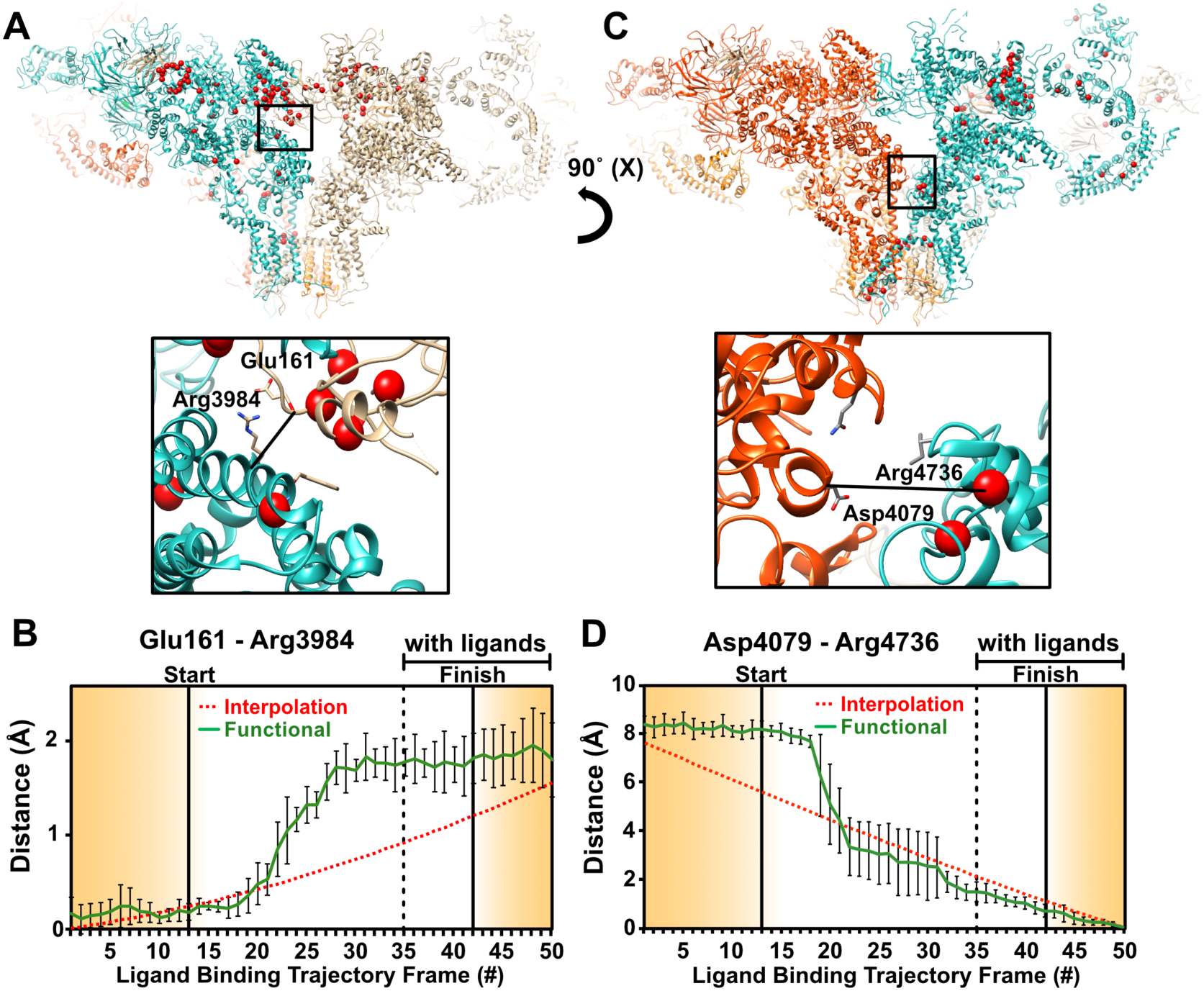
**RyR1 conformational changes at interprotomer contact sites along the functional trajectory**, augmented with excursions from the minimum-energy points along RC1. These excursions are in the increasing RC1 direction on the no-ligand landscape, and in the decreasing RC1 direction on the with-ligand landscape. **(A,C)** The general regions and the specific sites (insets) examined in each region. The monomers are shown in different colors. Red markers highlight known point-mutations involved in malignant hyperthermia 1, and central core disease of muscle. Distance variations are between two amino acid backbones at contact sites between the monomers. The amino acids used for measurement are shown as sticks. **(B, D)** Distance variations between carbon-alpha backbone atoms of two opposing residues for 50 structures along functional and interpolated trajectories. The measurements were performed and plotted as described in Fig. 6.

The second significant contact is situated between the NTD β8– β9 loop and the activation domain. These two domains move apart by ~1.5Å in a stepwise fashion as the interlandscape transition point is approached (Fig. 9A). The loop, containing a number of charged residues suitable for ionic interactions, has been suggested to play an important role in gating (*24*, *26*). This contact is therefore a “gate” candidate, where strong ionic interactions would prevent pore-opening before ligand binding. The contact would be broken by conformational changes in the activation domain upon ligand binding, thus allowing pore opening. Other, weaker “zipper” interactions would then be broken and reformed at intervals by the Brownian motion of the channel, accounting for the observed “flickering” behavior of RyR1. This tempting hypothesis for the RyR1 gating mechanism remains to be experimentally tested.

Finally, the IP3 receptor, with a homologous calcium activation mechanism, does not have EF-hands and an S2S3 domain, but its N-terminal domains are homologous to the RyR1 NTDs, where the activating ligand IP3 binding site is located (*30*, *31*). The IP3 receptor could thus have a homologous mechanism, where binding of calcium and IP3 lead to conformational changes in the activation domain and N-terminal domains of IP3R, followed by pore opening.

## Discussion

We now turn to the more general implications of our work. Inferring biological function from structure is a paramount goal of structural biology. Singly and together, the results presented here highlight the importance of basing functional inference on experimentally determined energy landscapes.

Our approach is based on three concepts with wide-ranging implications. First, biological function involves a rich set of *continuous* conformational changes inadequately described by discrete structures of unknown relationship. Second, *thermal fluctuations in equilibrium* lead to sightings of all states up to an energy limit set by the number of snapshots in the dataset. This makes it possible to compile the energy landscapes needed for a rigorous description of the thermodynamics of function. And third, the course of a biological process can be inferred by pooling data from ensembles in equilibrium with reservoirs corresponding to the initial and final states of the process, provided continuous, functionally relevant conformational changes can be mapped. We believe the energy landscapes, the inter-landscape transition maps, the new information on the conformationally active structural domains, and the nature, sequence, extent of important displacements involved in function – in this case ligand binding, channel activation, and channel gating – demonstrate the importance of the function-based analytical approach used here.

Clearly, equilibrium measurements cannot answer all questions regarding dynamics. It would thus be illuminating to compare the present results with those obtained from the observation of non-equilibrium ensembles engaged in reaction. Also, using larger equilibrium datasets, it would be interesting to investigate the role of conformations lying at higher energies. These constitute future tasks.

## Conclusions

We have presented a new approach to determining conformational changes associated with biological function. The results demonstrate the possibility of a rigorous description of biological function at high spatial resolution. The new insights include the nature, sequence, and extent of conformational motions involved in function, and the way allosteric signals are transduced to remote sites. The approach is general, and thus applicable to a wide range of systems and processes.

## Acknowledgments

We acknowledge valuable discussions with E. Lattman, G. Phillips, M. Schmidt, and members of the UWM data analysis group. The research conducted at UWM was supported by the US Department of Energy, Office of Science, Basic Energy Sciences under award DE-SC0002164 (algorithm design and development, and data analysis), by the US National Science Foundation under awards STC 1231306 (numerical trial models) and 1551489 (underlying analytical models), and by the UWM Research Growth Initiative. The work performed by JF was supported by HHMI, NIH GM55440, and NIH GM29169. The work performed by DBH and AG was supported by CUNY.

## Movie captions

**Movie 1: Evolution of the atomic model of RyR1 along the functional trajectory** of Fig. 1. Color bar shows the magnitude of the atomic displacements.

**Movie 2: Evolution of the atomic model of RyR1 obtained by linear interpolation** between the “START” and “FINISH” discrete structures (boxed classes in Fig. 2). Color bar shows the magnitude of the atomic displacements.

**Movie 3: Conformational evolution (backbone representation) in the pore domain along the functional trajectory** of Fig. 1. Color bar shows the magnitude of the atomic displacements. Singular value decomposition (SVD) was applied to the movie of atomic models to reduce noise.

**Movie 4: Conformational evolution (backbone representation) in the pore domain obtained by linear interpolation between two discrete structures.** Color bar shows the magnitude of the atomic displacements.

**Movie 5**: **Conformational evolution in the asymmetric unit of the activation core domain along the functional trajectory** of Fig. 1. Colored spheres indicate ligand-binding sites (yellow: Ca^2+^; magenta: caffeine; brown: ATP). Color bar shows the magnitude of the atomic displacements. Singular value decomposition (SVD) was applied to the movie of atomic models to reduce noise.

**Movie 6: Conformational evolution in the asymmetric unit of the activation core domain obtained by linear interpolation.** Colored spheres indicate ligand-binding sites (yellow: Ca^2+^; magenta: caffeine; brown: ATP). Color bar shows the magnitude of the atomic displacements.

## Supplementary Materials

### 1. Upper limit on the energy of accessible conformational states

The difference in the Gibbs free energy Δ*G* between two states with populations *N*_*A*_ and *N*_*B*_ is given by the Boltzmann relation, viz. *N*_*B*_ / *N*_*A*_ = exp (−Δ*G / k_B_T*), with *k*_*B*_ the Boltzmann constant, and *T* the absolute temperature. Using an ensemble of *N* snapshots, the state sighted only once lies at an energy Δ*G*_*max*_ above the lowest energy state of the ensemble. Under these conditions, *N*_*B*_ = 1 and *N*_*A*_ = *N* −1 ≈ *N*. Thus, the highest energy observed in an ensemble of *N* particles corresponds to Δ*G*_*max*_ = *k_B_T* log *N*. For the dataset analyzed in this paper, *N*_*-ligand*_=293,619 and *N*_*+ligand*_=262,022, yielding a theoretical upper limit Δ*G_max_* ~ 7 kcal/mol.

The above approach for deducing free energies used, e.g., in (*1*, *32*) and elsewhere, relies on a number of assumptions. From Maxwell-Boltzmann statistics of non-interacting particles in thermal equilibrium, the occupation probability of a discrete state *i* is given by:

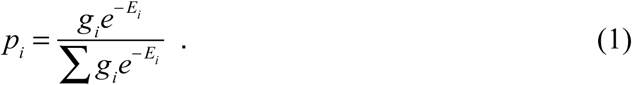

*g*_*i*_ is the degeneracy of the state *i*, specifically, the number of experimentally indistinguishable conformational states assigned to energy *E*_*i*_, which may nevertheless be distinguished from each other by some other means. *E* = *U*, *H*, or *G* (internal, Helmholtz, or Gibbs free energy, respectively). The sum extends over all possible states.

Conformational sorting yields the number of sightings (snapshots) of each conformation. Each sighting represents a conformational state occupied in thermal equilibrium. A conformational bin contains all conformations *indistinguishable by the experimental and data-analytical pipeline used* (“coarse-graining”).

The experimental observable, namely the number of sightings of a conformational state *c* is given by:

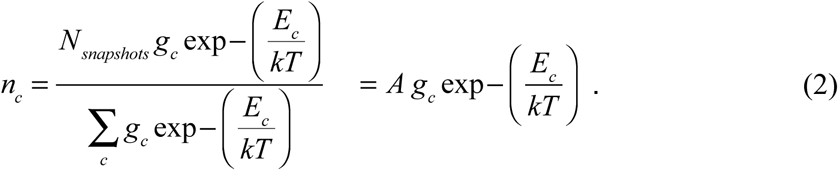

In principle, Eq.(2) provides a direct link between the number of sightings (snapshots) of a conformation, and its energy.

However, the degeneracy *g*_*c*_ induced by coarse-graining is unknown, and, in general, conformation-dependent. *g*_*c*_ can be absorbed into the exponent via an entropy term, viz.

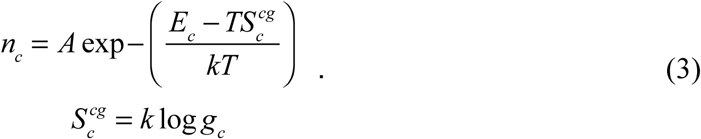

with the superscript *cg* short for coarse-graining.

Eqs. (2) and (3) implicitly assume that all conformations coarse-grained into the same conformational class have the same energy. This need not be the case. Without further information or assumptions, it is not possible to determine energy differences between conformational classes from Eqs. (2) or (3).

Mapping conformational landscapes without regard to the degeneracy induced by coarse-graining (as commonly practiced) is predicated on the further assumption that the variance of *E*_*c*_ dominates over that of 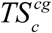, viz.
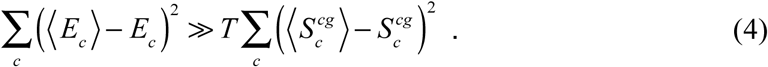

In this work, Eq. (3) is used subject to the assumption that Eq.(4) holds.

### 2. Input data, preprocessing steps, and analytical pipeline

The details of the data used for the present analysis have been described elsewhere (*10,12*). Here, we provide a brief outline for the reader’s convenience.

#### Rabbit Skeletal Muscle RyR1 Purification

Purified RyR1 was prepared as described in (*10*). Snap-frozen rabbit skeletal muscle was blended in cold buffer containing 10mM Tris-Maleate pH-6.8, 1mM DTT, 1mM EDTA, 150μM PMSF and 1mM Benzamidine, and centrifuged for 10 minutes at 8000g. The supernatant was centrifuged for 20 minutes at 40,000g. Pellets were solubilized in 10mM HEPES pH=7.5, 1% CHAPS, 1M NaCl, 2mM EGTA, 2mM TCEP, and protease inhibitors cocktail (Roche). The solubilized membranes were then diluted 1:1 in the same buffer without the NaCl, and centrifuged for 30 minutes at 100,000g. The supernatant was then passed through a 0.2 micron filter and allowed to bind overnight at 4 °C to a pre-equilibrated 5ml GStrap (GE healthcare) column with bound GST-Calstabin1. The column was then washed with modified solubilization buffer (0.5% CHAPS and 0.5M NaCl), and RyR1 was eluted with two column volumes of 10μM Calstabin2 in the same buffer. RyR1 was then concentrated on a 100,000kDa cut-off centrifugation filter and run through a size exclusion column (tosoh G4SWxl). The mono-disperse peak was then concentrated on a 100,000kDa cut-off centrifugation filter to ~5-10mg/ml.

#### Residual Ca^2+^ concentration in the no-ligands solution

The buffer (0.5M NaCl) was potentially contaminated with 0.002% Ca^2+^, corresponding to 10µM Ca^2+^ in solution. 5mM of EGTA was added to chelate the solution. A small concentration of Ca^2+^ ions remains in the no-ligands solution after such treatment. According to http://maxchelator.stanford.edu/CaMgATPEGTA-TS.htm, this free Ca^2+^ concentration is 0.2nM. Similar concentrations are reported in: http://onlinelibrary.wiley.com/doi/10.1113/jphysiol.1968.sp008413/epdf

We now investigate whether binding of residual Ca^2+^ contamination is responsible for displacing the RyR1 molecules from the no-ligand minimum-energy region to functionally significant, higher-energy regions of the no-ligands landscape.

- Ca^2+^ contamination in “no-ligands” solution: 2×10^−10^M
- Total RyR1 in “no-ligands” solution (5mg/ml, 2.3MDa): 2×10^−6^M
- Number of RyR1 at no-ligands transition point (2% of total): 4×10^−8^M
- Assuming all Ca^2+^ is bound, the ratio [No. of Ca^2+^-contaminated RyR1 molecules] / [No. of RyR1 molecules at transition point] is: 2×10^−10^/ 4×10^−8^ = 5×10^−3^ =0.5%

The concentration of Ca^2+^ contaminants is ~ 0.5% of the number of RyR1 molecules in functionally important, high-energy regions of the landscape, such as the inter-landscape transition points. The role of any Ca^2+^ contamination is thus negligible.

#### Cryo-Electron Microscopy

The RyR-EGTA and RyR-30µM Ca^2+^-ATP-caffeine samples were prepared on holey carbon grids (C-flat CF-1.2/1.3-2C-T, Protochips Inc, NC), as described previously (*10,12*). 3 µL of each sample was applied to holey-gold grids, blotted for 3.5 to 4 seconds and vitrified by rapidly plunging into liquid ethane with a Vitrobot (FEI). Data were acquired using an FEI Tecnai F30 Polara (FEI, Eindhoven) operating at 300 kV with the automated data collection software Leginon (*33*) on a K2 Summit direct electron detector camera (Gatan, Pleasanton, CA) at a nominal magnification of 31,000x, corresponding to a calibrated pixel size of 1.255 Å. For experiments using carbon grids, images were recorded in dose-fractionated mode, each image being fractionated into 20 frames. The total exposure time was 4 s, yielding a total accumulated dose of 25 electrons/Å^2^ on the specimen. The beam diameter was set at approximately 500 nm in order to capture two images per hole using image shift.

#### Image Processing

Dose-fractionated image-stacks collected with a Gatan K2 Summit camera were aligned using MotionCorr (*34*) and the sum of aligned frames was used for further preprocessing. The particles were picked with RELION 1.3 (*5*) reference-based automated particle picking procedure (*35*) and their defocus values estimated by ctffind4 (*36*). Particles were subjected to 3D classification using RELION 1.3 to select the good particles. In this way, 366,000 particles from the ligand-free dataset and 450,000 particles from the ligand-bound dataset were selected.

#### Orientation Recovery

The orientation parameters for each of the two datasets were then refined separately using RELION 1.3 with the same starting reference and no symmetry imposed. These alignment parameters were subsequently used for the manifold-based analysis, as described in (*1*, *37*). In brief, aligned, centered snapshots were divided into projection directions, specifically, into groups falling onto tessellations of a spherical shell subtending a semi-cone angle of two Shannon angles (defined as spatial resolution (0.4 nm) / particle diameter (32 nm)). Embedding the snapshots in each projection direction by Diffusion Map (*38*) revealed two clusters, one corresponding to an “artifact class” of unusually low contrast. This class was excluded, leaving 791,956 snapshots for further analysis.

The geometric machine-learning analytical pipeline, described in detail in (*1*), consists of the following steps:

a. Diffusion Map embedding of ±ligand snapshots in each projection direction.
b. Nonlinear Laplacian Spectrum Analysis (NLSA) (*12*) after 100-fold concatenation along each of the first two eigenfunctions of the Laplace-Beltrami operator.
c. Compilation of the energy landscapes of the ±ligand datasets.
d. Mapping of the inter-landscape transition probability, using the formalism described in section 3 below.
e. Compilation of 3D movies along selected trajectories on the energy landscapes.

### 3. Estimating transition probabilities by Fermi’s Golden Rule

Fermi’s Golden Rule (*15*, *16*) provides the following expression for the (non-equilibrium) transition rate between an occupied initial state *i* and a final state *f*:

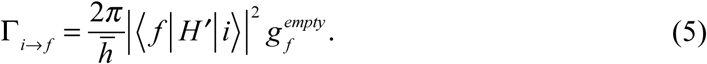

Γ_*i*→*f*_ is the transition rate between *i* and *f*, 〈*f*|*H*′|*i*〉 the matrix element between the initial and final states, 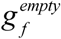 the number of unoccupied final states per energy interval (“density of states”), and 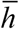 the Planck constant divided by 2π.

For a ligand-binding transition between two conformations *c*, *c*′, Eq.(5) becomes:

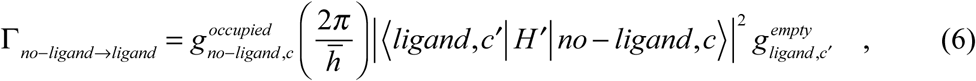

where 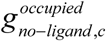 is the number of occupied initial states per energy interval.

Consider an ensemble of entities without ligand (but capable of binding ligands) in equilibrium with a thermal reservoir. The introduction of a reservoir of ligands induces the ensemble to evolve toward a new equilibrium, in which the fraction of +ligand entities no longer changes with time.

Consider the initial stages of the approach to equilibrium with the ligand reservoir. At times short compared with the ensemble relaxation time, the occupied conformational spectrum is nearly the same as that prior to contact with the ligand reservoir. Under these conditions, the number of occupied conformational states 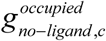 can be deduced from Eq.(2), viz.

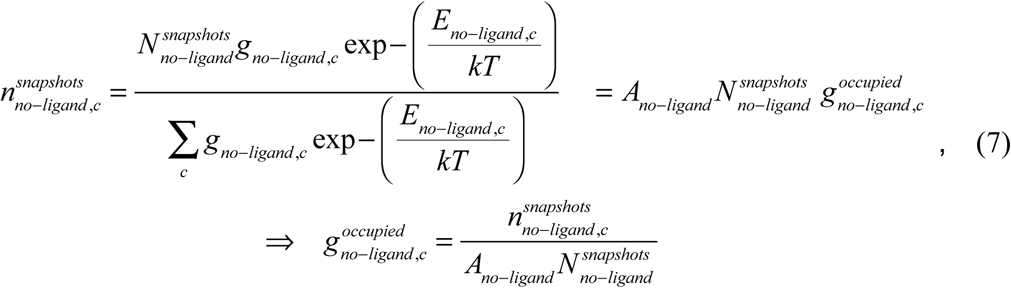

with 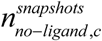 representing the number of snapshots per energy interval from the no-ligand ensemble in conformational bin *c* and 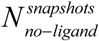 the total number of snapshots from the no-ligand ensemble.

In the early stage of relaxation considered here, essentially all +ligand states are empty. Eq.(7) can then be used to determine 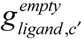 as follows:

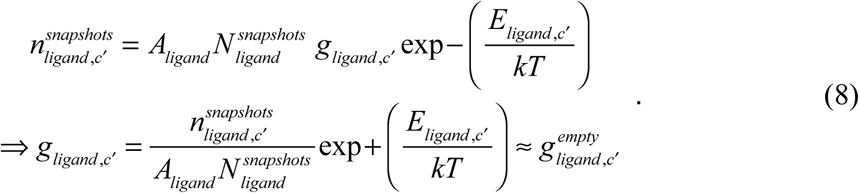

Eqs.(7) and (8) provide the *g*-factors in Eq.(6), for times short compared with the ensemble relaxation time.

In general, the matrix element is unknown. Here, we approximate the square-modulus of the matrix element with an appropriately scaled delta function, viz. *E*′δ(*c,c*′), with *E*′ an energy corresponding to the perturbing Hamiltonian *H*′. This assumes ligand binding connects iso-conformational states on the −ligand and +ligand landscapes. In this model, the inter-landscape transition *per se* is too rapid to allow conformational adjustments, but may be predicated on prior conformational changes, and/or initiate subsequent conformational adjustments. The expression for the transition rate per energy interval now becomes:

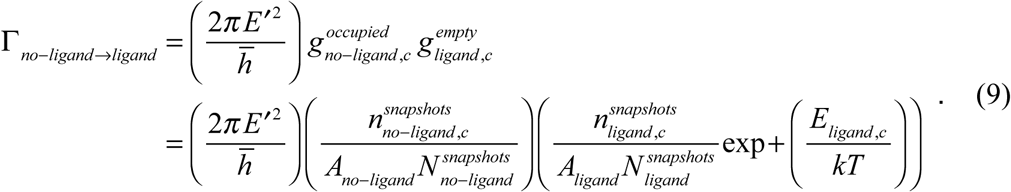

The ratio of transition rates at two points on the conformational landscape is given by:

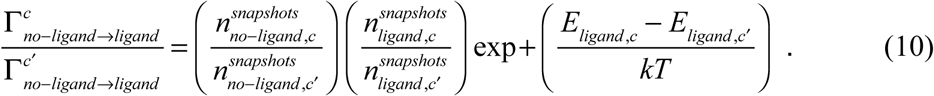

Eqs.(9) and (10) become inaccurate, however, when the number of snapshots per conformational bin is dominated by Poisson noise.

### 4. Nature of conformational changes along reaction coordinates

NLSA (singular value decomposition on the manifold) (*12*) with the data ordered according to their projection on a given reaction coordinate reveals the evolution of the data along that reaction coordinate. Using the snapshots in one projection direction, Fig. S1 shows the conformational deviation of the data from the mean along RC1 and RC2.

### 5. Estimating the spatial resolution of the density maps

The procedure of comparing independent half-set reconstructions via Fourier shell correlation is known as the “gold standard” in resolution estimation. This approach cannot be readily used to estimate the resolution of our maps, because division of data into two subsets at the outset reduces the conformational sampling to a level incompatible with reliable analysis. This is already a limiting factor in our analysis, as evidenced by the need to define an orientational aperture radius four times the size commensurate with the resolution of the data of 0.4 nm (*12*) as estimated by RELION. Division of the data at later points along the analytical pipeline does not produce independent datasets.

We therefore use two alternative means to estimate the resolution of our density maps, as follows.

a. The program ResMap (*39*) estimates the local resolution as ranging from 0.35 nm in the core to 1.2 nm at the outer edge of the C4 symmetrized maps.
b. Structural features in the map core evidently correspond to a resolution of approximately 0.4 nm, with bulky side chains visible in the best parts of the map (Fig. S2). Resolution in the outer parts of the map is much lower, around 1.2 nm, in large part due to the coarse angular sampling of the data, which limits the resolution in a radius-dependent manner. The original RELION analysis reached ~ 0.4 nm in the core, similar to the resolution observed here on the innermost parts of the molecule.

### 6. Fitting and refinement of atomic coordinates

Each of the 50 maps along the trajectory was fitted to a model with the rigid-body fit facility in COOT (*40*), using multiple starting models to avoid model bias (PDB ID: 5TB4, 5T9R, 5TAP, 5T9V, 5TAL, 5TAQ) (*9*). The models were then refined in real-space using phenix.real_space_refine (*41*).

Distance measurements between residue pairs for each of the 50 maps were performed with UCSF Chimera (*42*) using residue backbones as references. The distances obtained from different starting models were then averaged. The error bars show the full scatter (not standard deviation) of the results obtained with different starting models.

The models were fitted to each 50 maps along the trajectory using COOT (*40*), starting from the model of the Ca^2+^-ATP-Caffeine open state (PDB ID: 5TAL) (*9*), rigid-body fit into each map. The models were then refined in real space using phenix.real_space_refine (*41*).

Distance measurements between residue pairs for each of the 50 maps were performed with UCSF Chimera (*42*) using residue backbones as references.

### 7. Nature of conformational changes along a ligand-binding trajectory

Maps and models built along the trajectory show domain motions such as up-down shell movement, rotation of the activation domain, pore opening and rotation of the pseudo-voltage sensor domain (pVSD). The concerted nature of these motions may be the result of direct coupling between the respective domains (Fig. 9 & Fig. S3 and S4).

### 8. Computational Resources

All computations were performed on a CPU cluster with following specifications: 16 CPU nodes, each consisting of two Deca-core E5-2660 V3 “Haswell”/2.6GHz and 128GB of memory. Particle picking, contrast transfer function (CTF) estimation, and initial orientation recovery were performed using RELION. 3D back-projection on 2D NLSA snapshots was executed using the reconstruct function in RELION. UCSF Chimera was used for visualization and compilation of 3D movies (*42*).

### 9. Data availability

The density maps and the atomic models along the transition trajectory will be deposited in EMDB and the Protein Data Bank, respectively.

---

**Fig. S1:**
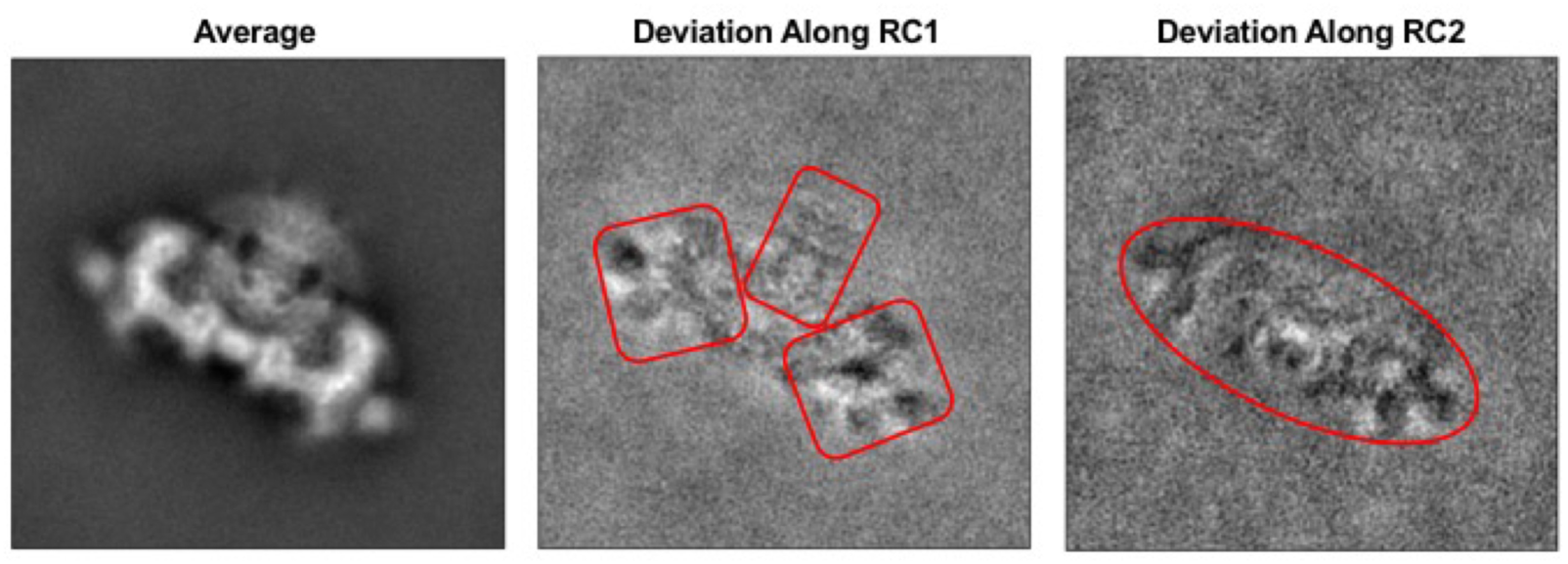
**Average structure and deviations from the average structure of RyR1 along Conformational reaction coordinates (RC) 1 & 2**, respectively, viewed in one projection direction. The deviations from the mean reveal the concerted conformational changes associated with each reaction coordinate.

**Fig. S2:**
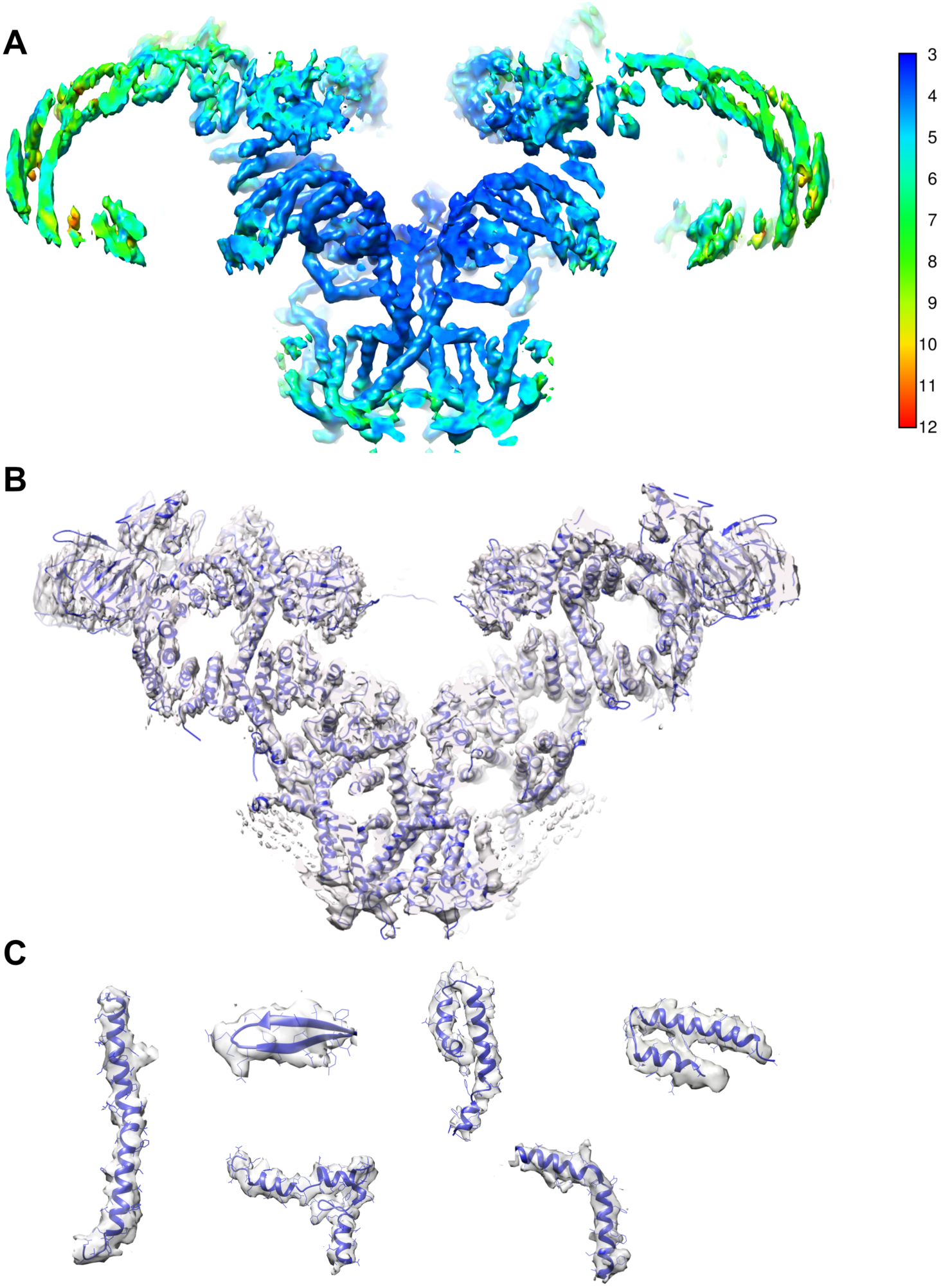
Estimate of spatial resolution. **(A)** Slab through one of the cryo-EM reconstructions of RyR1 along the trajectory, colored according to the local resolution as measured by ResMap. **(B)** Slab through the same cryo-EM map fitted with the atomic model of RyR1. Map and model are shown in transparent surface and ribbon representations, respectively. **(C)** Densities and models of subsets of secondary structure elements in different regions of the map.

**Fig. S3:**
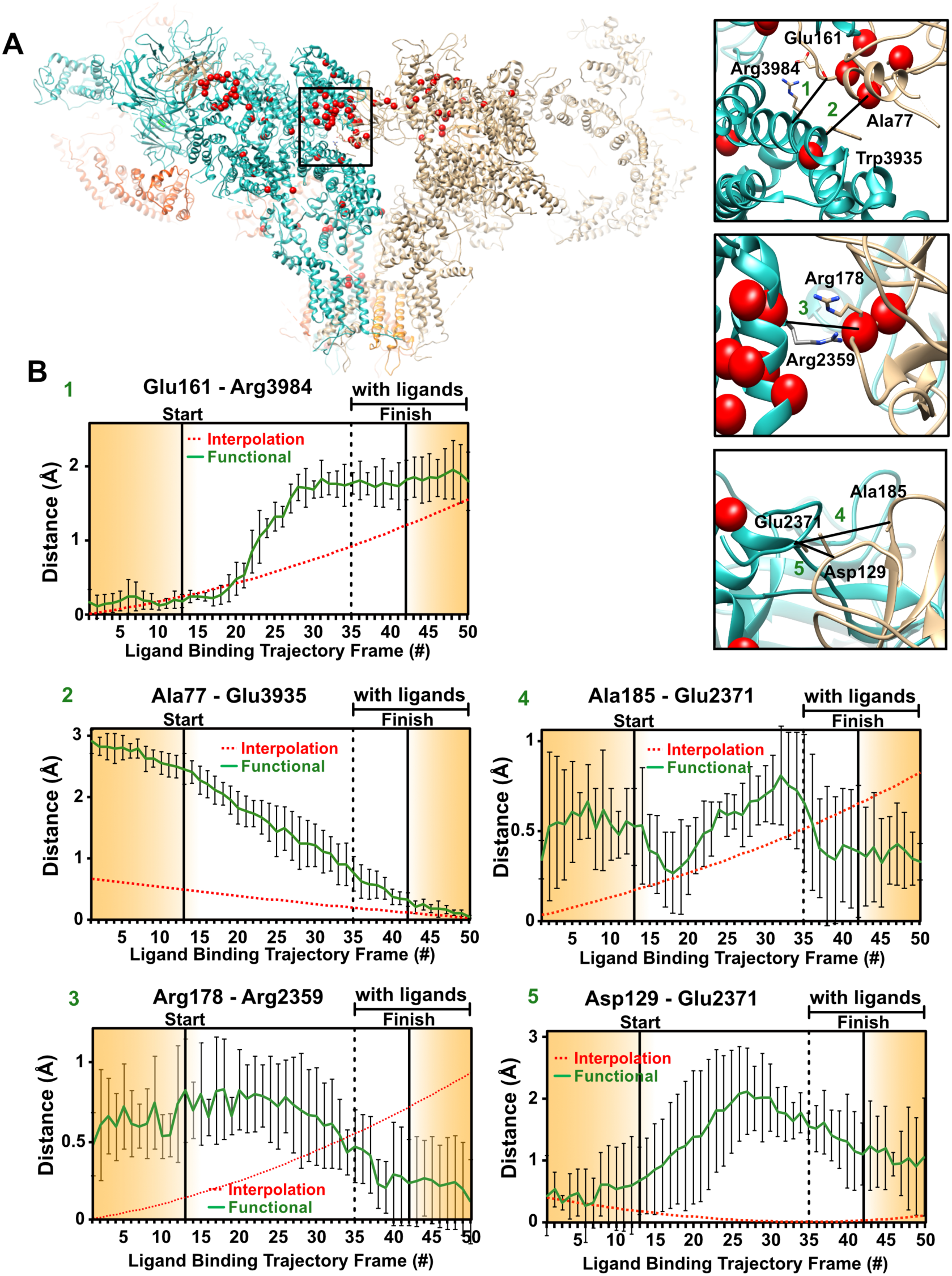
**RyR1 changes at interprotomer contact sites in N-terminal domain along the route of Fig. 1(A)**, augmented with excursions from the minimum-energy points along RC1. These excursions are in the increasing RC1 direction on the no-ligand landscape, and in the decreasing RC1 direction on the with-ligand landscape. **(A)** The general region, and the specific sites examined in detail, with each monomer shown in different colors. The locations of known point-mutations found in malignant hyperthermia 1 and central core disease of muscle are shown in red. **(B)** Distance variations measured between carbon-alpha backbone atoms of two opposing residues for each of 50 frames along the functional trajectory. The measurements were calculated and plotted as described in Fig. 6. Each measured distance between two amino acid backbones at contact sites between the monomers. The amino acids used for measurement are represented in sticks.

**Fig. S4:**
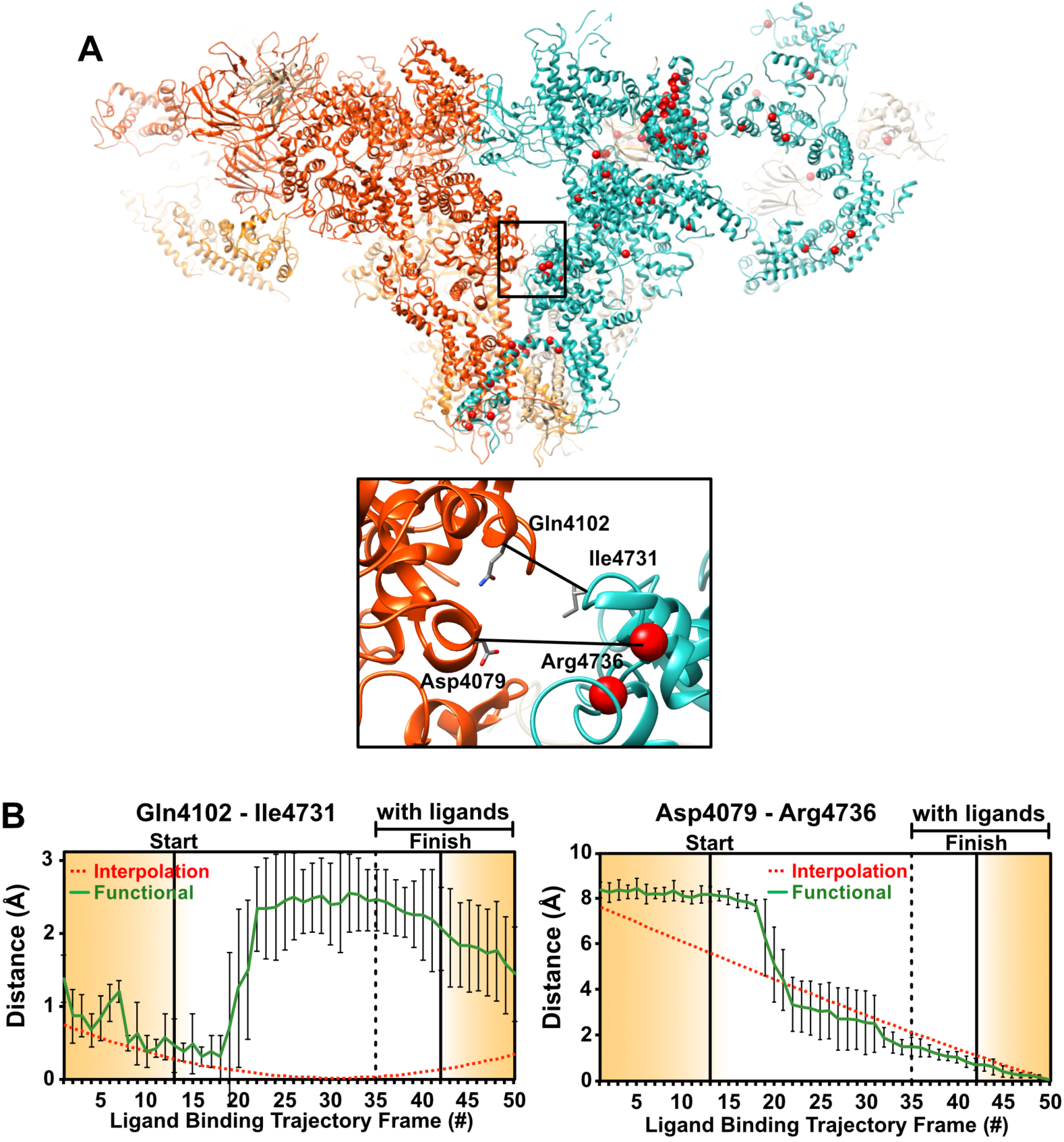
**RyR1 changes at interprotomer contact sites in activation core along the route of Fig. 1(A)**, augmented with excursions from the minimum-energy points along RC1. These excursions are in the increasing RC1 direction on the no-ligand landscape, and in the decreasing RC1 direction on the with-ligand landscape. **(A)** The general region, and the specific sites examined in detail, with each monomer shown in different colors. The locations of known point-mutations found in malignant hyperthermia 1 and central core disease of muscle are shown in red. **(B)** Distance variations measured between carbon-alpha backbone atoms of two opposing residues for each of 50 frames along the functional trajectory. The measurements were calculated and plotted as described in Fig. 6. Each measured distance between two amino acid backbones at contact sites between the monomers. The amino acids used for measurement are represented in sticks.

## Movie Captions

**Movie S1:** Conformational changes along the functional trajectory of Fig. 1 (trans-membrane view).

**Movie S2:** Conformational changes along the functional trajectory of Fig. 1 (cytoplasmic view).

**Movie S3:** Conformational changes along the functional trajectory of Fig. 1 obtained by modeling (trans-membrane view).

**Movie S4:** Conformational changes along the functional trajectory of Fig. 1 obtained by modeling (cytoplasmic view).

## Author Contributions

AD designed and implemented the data-analytical approach, a robust pipeline for driving energy landscape, transition probability maps and continuous conformational changes from cryo-EM snapshots.

AG processed the cryo-EM data using standard methods. AG and DBH designed the molecular modeling and distance measurement strategy. DBH performed the molecular modeling and distance measurements.

GM helped with implementation of data-analysis algorithms, mainly for the energy landscapes and the continuous conformational movies.

PS co-developed the geometric manifold-based approach, co-investigated the effect of coarse graining on energy landscapes, and helped implement the software.

JF co-designed the study, analyzed the results in terms of the concepts of single-particle cryo-EM and the dynamics of molecular machines, and co-wrote the paper.

AO co-designed the study and the manifold-based data-analytical algorithms, identified and co-analyzed the effect of coarse-graining, proposed the ligand-binding model and formalism for estimating interlandscape transition probabilities, and wrote the paper.

AO, AD, AG, DBH and JF analyzed and interpreted the results.

